# Host usage in *Aedes aegypti* collected from Houston, Texas and Phoenix, Arizona using whole mosquito third-generation sequencing blood meal analysis

**DOI:** 10.1101/2025.05.19.654806

**Authors:** Brittani A. Ciomperlik, Edwin R. Burgess, Neil D. Sanscrainte, Mba-tihssommah Mosore, John Townsend, James Will, Nicole Busser, Alden S. Estep

## Abstract

*Aedes aegypti* is the primary vector of several viruses of international public health concern, including Zika, dengue, yellow fever, and chikungunya. Their synanthropic ecology and establishment in tropical, sub-tropical, and temperate areas make *Ae. aegypti* one of the most medically relevant mosquito species. While they have been reported to be highly anthropophilic, several studies indicate a broader host range. They are also reported to take multiple blood meals between gonotrophic cycles. Consumption of multiple blood meals makes determination of host usage difficult when using common blood meal analysis methods. In this study, we examined host usage of *Ae. aegypti* in Harris County, Texas (Houston), and Maricopa County, Arizona (Phoenix), using a Nanopore-based third-generation sequencing protocol to successfully resolve host usage multiplicity and identify a broad range of host usage. Using this method, approximately 80% of samples from each location with evidence of blood feeding resulted in a blood meal species identification, with a single host blood meal in about 80% of samples and approximately 20% containing evidence of multiple blood meals. Overall, we observed a wide host range with human DNA the most prevalent followed by feline and canine. We also identified avian, rodent, ungulate and even ectotherm usage by *Ae. aegypti* from Maricopa County. The discovery of hosts other than humans expands the understanding of host usage dynamics in *Ae. aegypti* and their involvement in arbovirus transmission systems in highly urbanized areas such as Harris and Maricopa Counties.

**Synopsis:** The yellow fever mosquito, *Aedes aegypti*, is a vector of several viral pathogens that frequently cause human disease in tropical and subtropical locations but many of these pathogens are not controlled by vaccination. As such, control of this mosquito is critical to reduce disease transmission and better understanding of the ecology of this species helps to improve the efficacy of operational control. In this study, researchers used a deep sequencing method to define patterns of host usage in two large US metropolitan areas, Houston, TX and Phoenix, AZ, with substantial *Ae. aegypti* infestations. Results identified frequent multiple blood feeding by individual mosquitoes and that the primary host sources were humans and their pets. Results from Phoenix indicated a broad host usage, including feeding on livestock, birds, and even reptiles. This may provide useful information about *Ae. aegypti* resting sites and could be used to improve treatment by operational personnel.

## Introduction

Both host usage and host range are key factors in cycles of mosquito borne pathogen transmission [1]. Host usage, or host utilization, is defined as the preference for certain species and the frequency a certain host is used [2], while host range is the behavioral plasticity to shift from a preferred host to an available one, expanding the number of species, genera, or families utilized [1, 3]. Pathogen transmission cycles are better understood when there is adequate knowledge of the identity and frequency of hosts that are chosen by mosquitoes for blood meals [4, 5]. Blood meal analysis (BMA) assists in reaching a comprehensive understanding of what hosts are being utilized and, with the use of third-generation sequencing (TGS), a deeper approach can be taken to monitor vector-to-host contact rates as well as pathogen transmission.

Blood meal analysis received attention after Reeves et al. [6] described the zoonotic relationship between vertebrate hosts (i.e. human or domesticated animals) with regards to some arboviruses. For decades, most BMA studies were conducted using several antibody-based methods that required specific antibodies for each host or host family being assessed [7–13]. Antibody based methods are sensitive but necessarily limited in host identification by the range of host specific antibodies available; to identify usage of a host, it requires the foreknowledge to include an antibody to detect the host. Currently, an increasingly common BMA method involves the selective amplification of host DNA by polymerase chain reaction (PCR) with specific primers and then sequencing of the amplicon using the Sanger method [2, 14–18]. The primers used for BMA are designed to amplify DNA from a taxonomic spectrum of potential hosts and limit co-amplification of non-target DNA [16]. These DNA based methods increase the possibility of identifying hosts since results are compared to large sequence databases. DNA based BMA has increased the understanding and range of host usage and often allows identification to the species level. Limitations using the Sanger method include its inability to differentiate multiple blood meals taken by a single mosquito without extensive additional efforts (i.e. cloning to isolate individual host sequences), the potential for co-amplification of the mosquito DNA with general primers, and the limited number of host-specific primer sets currently available for the *cytochrome c oxidase subunit I* (*COI*) gene commonly used in reference databases as well as the scope and quality of *COI* databases [16].

Next generation and TGS based methods have been recently developed for BMA and overcome some of the limitations of Sanger sequencing. Like any method based on sequence comparison, a strict quality filtering process is needed to ensure proper assignment of reads to both hosts and samples. These methods, like Sanger based methods, cannot overcome issues with *COI* databases, but the ability to assign individual reads from the same amplified sample to different hosts is the most notable benefit and moves host identification beyond that possible with Sanger and antibody based methods [19, 20].

*Aedes aegypti* is the primary vector of Zika, dengue, yellow fever, and chikungunya viruses, making it one of the most medically important mosquitoes [12]. It is an important vector of pathogens in subtropical and tropical countries around the globe. The adaptivity of *Ae. aegypti* has enabled it to go from a primarily sylvatic species, with access to a wide range of hosts, to a more urban one, breeding in artificial containers of fresh water, which are abundant in human inhabited areas [21]. Studies of the feeding behavior of *Ae. aegypti* have demonstrated that it is anthropophilic but has significant plasticity in host usage (reviewed in Olson et al. [22]).

Previous studies on *Ae. aegypti* often identify feline and canine hosts in addition to human, and less commonly other mammalian and avian hosts [9, 17, 22–24]. Studies of *Ae. aegypti formosus*, considered the original sylvatic form, have the wide host range expected of a mosquito endemic where humans are infrequent [9, 20]. A similarly broad host range has been shown in the more domesticated form of *Ae. aegypti* from northern Mexico, which identified the presence of reptile blood [25]. Host preference, and the components which determine that preference, play an important role in how diseases are transmitted to humans by *Ae. aegypti* [1].

The variability of host usage and host range found in these previous studies, combined with limited studies in *Ae. aegypti* found in the US, led us to the present study [17, 18, 22, 26]. Our goal was to describe host usage of blood fed *Ae. aegypti* collected from Harris County, Texas, and Maricopa County, Arizona; both are large metropolitan areas (Houston, TX and Phoenix, AZ) with extensive infestations of *Ae. aegypti* and no published information about host usage in *Ae. aegypti*. Our objectives were two-fold. First, to develop a sensitive but time and resource efficient Nanopore TGS method that could detect blood meals in the presence of substantial mosquito DNA, then to utilize this method to determine host usage, host range, and frequency of host multiplicity of *Ae. aegypti* from normal vector control surveillance collections. This study expands our understanding of *Ae. aegypti* host usage in the US and provides researchers with a novel and flexible tool to enhance blood meal identification capabilities in mosquitoes.

## Materials and Methods

### Sample collection

Mosquitoes were collected in Harris County, TX using BG-Sentinel traps (Biogents AG, Regensburg, Germany) by Harris County Mosquito and Vector Control Division (MVC Division) employees from June-September of 2019. Samples were preserved on ice in the field and then identified morphologically to species while under cold anesthetization at the MVC Division. Samples were stored at -80 °C and then shipped frozen to the USDA-ARS-CMAVE where *Ae. aegyp*ti pools were separated and homogenized individually for standardized insecticide resistance genotyping [27, 28]. Sample homogenates that had an orange or red hue, indicative of the presence of a blood meal, were refrozen at -80 °C for use in this study (Figure 1, Step 1). A total of 31 of the 440 mosquito homogenates appeared positive for blood meal presence. Three mosquitoes with no evidence of a blood meal were extracted to be used as non-blood fed controls.

**Figure 1:**
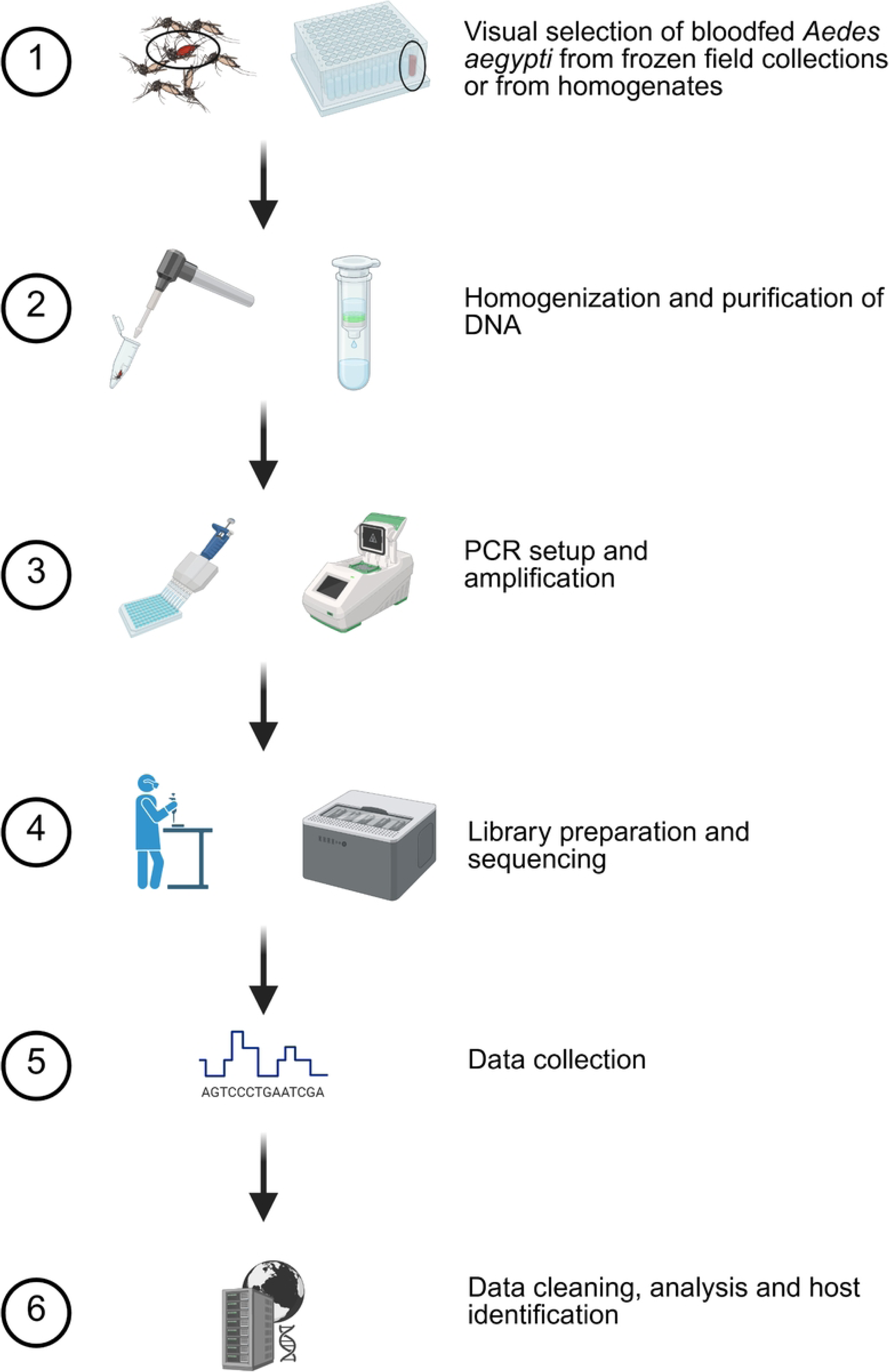
Blood meal analysis methodology. Based on preliminary studies, a six step procedure was used to determine host usage. Mosquitoes with evidence of blood were selected for homogenization and purification. Specific details are provided in the methods section. Created in BioRender (https://BioRender.com/m4ubsfn).

Mosquitoes from the Phoenix, AZ metropolitan area were collected using encephalitis vector surveillance/carbon dioxide baited traps or BG-Sentinel traps from May-October 2021 and then identified to species by Maricopa County Environmental Services Department personnel [29, 30]. Traps were set ∼1:00 PM, allowed to run overnight, and collected ∼10:00 AM the next morning. Frozen *Ae. aegypti* were shipped to USDA-ARS-CMAVE for assessment of genetic insecticide resistance markers. Approximately 5,000 *Ae. aegypti* were screened for the visible presence of a blood meal (Figure 1, Step 1) and these blood fed organisms were stored at -80 °C until processing.

### Preliminary method development

Preliminary studies were conducted to determine an optimal amplification temperature, to pilot the use of unpurified PCR products as direct input into the nanopore-based sequencing library preparation procedure, and to define quality control procedures and thresholds to ensure accurate assignment of reads. Details on these studies are in supplemental information (Supplemental File 1). During the course of this study, a nanopore-based method for host identification was published by Kothera [31] for *Culex* mosquitoes.

### Nucleic acid purification and amplification

DNA from the Harris County samples was purified using a silica column kit (gDNA kit, Zymo Research, Irvine, CA) from homogenates (300 μL) diluted in lysis buffer (1,200 μL) and processed per the manufacturer’s protocol for liquid samples. Maricopa County *Ae. aegypti* were manually homogenized in 350 μL of lysis buffer using a RNase-free disposable pellet pestle (ThermoFisher, Waltham. MA) or mechanically using 2.0 mm zirconia beads (BioSpec Products, Bartlesville, OK) (Figure 1, Step 2). DNA was purified using the Zymo Research Quick-DNA/RNA Miniprep Kit or the Zymo Research Quick-DNA Miniprep Kit following manufacturer’s instructions. DNA was eluted in 50 μL and frozen until PCR and sequencing.

*Cytochrome oxidase I* PCR was conducted using the primers, amplification conditions, and modified methods from Reeves et al. [16] (Figure 1, Step 3). Amplification was performed in 20 µL reactions using 2.0X Apex Taq RED Master Mix (Genesee Scientific Corp., San Diego, CA), primers VertCOI_7194_F 5’- CGMATRAAYAAYATRAGCTTCTGAY- 3’ (0.47 µM) and Mod_RepCOI_R 5’- TTCDGGRTGNCCRAARAATCA -3’ (0.47 µM), and 1 uL of DNA template at the following conditions: 95 °C for three minutes, followed by 40 cycles of 95 °C for 40 seconds, 50 °C for 30 seconds, and 72 °C for three minutes, with a final extension step of 72 °C for seven minutes. Amplicons were visualized on 1% agarose gels in tris-acetate EDTA buffer using GelRed (Biotium, Inc., Fremont, CA) alongside TrackIt 100 bp DNA Ladder (ThermoFisher, Waltham, MA) and imaged on an iBright system (ThermoFisher, Waltham, MA). The expected amplicon size, including the primers, is 440 bp.

### Nanopore sequencing

Based on successful preliminary Nanopore sequencing trials (see supplemental information), unpurified PCR amplicons were used with the R10 chemistry native barcoding kit 96 V14 (LSK-SQK-NBD114.96, protocol version: NBA_9170_v114_revL_15Sep2022).

MinKNOW software (versions 23.07.5 & 23.07.12) was used to manage sequencing and base calling on a GridION device (Oxford Nanopore Technologies, Oxford, UK). The manufacturer’s protocol was followed for sequencing sample preparation with one modification: 3 μL of unpurified PCR product was used as input into the end preparation step (rather than 50 femtomoles as directed by the manufacturer’s protocol) (Figure 1, Step 4). Samples were sequenced in batches of 96 samples per flow cell for a maximum 72-hour sequencing period and a minimum quality threshold of 10 was set in the MinKNOW software (Figure 1, Step 5). A working protocol for this methodology is provided in the data repository (DOI: 10.15482/USDA.ADC/26018947).

To ensure a rigorous quality filtering process that would reduce misassignment of barcodes by MinKNOW and account for the higher error rate of Nanopore sequencing (which could lead to misassignment of reads), we included several controls. Each individual flow cell contained two controls: one “blood negative” sample as a barcoded library from a non-blood fed laboratory *Ae. aegypti* and one “DNA negative” as a library that had only barcode but no DNA. To further ensure the process was sound, one 96 library preparation had 8 barcodes purposefully excluded to assess misassignment.

### Bioinformatic analysis

Bioinformatic analysis was conducted on a Dell Xeon laptop running Windows System for Linux using the FASTQ and sequencing summary files for each Nanopore sequencing run. The sequencing summary file was used for initial quality control and a custom script was used to remove low quality reads and ensure the presence of barcodes of at least 37 bases on each end of a read and a minimum 95% identity to the known barcode sequence. Following this initial quality control filtering, host identification was conducted by Basic Local Alignment Search Tool (BLAST version 2.6.0) [32] query against a custom database created from the Barcode of Life Database (BOLD) Systems [33, 34] that contained approximately 500,000 chordate *COI* sequences with species names. The resulting output files were filtered again to only include reads with a minimum 97.5% match and with an alignment length of at least 400 bases. This filtered file was then separated by barcode and blood meal identification. The minimum read threshold to be considered a species level match was two full length reads (>400bp) that met the above filters. The sequencing summary files, the BLAST formatted database, and the pipeline used in this analysis are available in the data repository. Raw sequencing data were submitted under Bioproject PRJNA1242810 and SRA accession numbers SAMN47605645-SAMN47605929.

### Statistical Analysis

Independence of single host frequency vs. multiple host frequency, as well as independence of feline, canine, and human host frequency between Harris and Maricopa counties, was analyzed using a Fisher’s Exact test in R version 4.4.2. [35]

## Results and Discussion

### Nanopore sequencing and quality control

Sequencing results indicated that the use of unpurified PCR amplicons resulted in successful output of data but, as expected, output was lower than when using purified amplicons. Rather than the manufacturer’s expected output of 10-20 Gbases per flow cell, we averaged 7.13 ± 3.20 Gbases and 12.15 ± 5.10 M reads per cell across three sequencing runs. Estimated N50 was consistent at 564.7 ± 1.2 bases. Approximately 84.6% of reads met the initial quality filtering by the MinKNOW software. Barcode binning was also managed by the software and 88.3 ± 2.5% of reads were binned. Sequencing run data is provided in the data repository (DOI: 10.15482/USDA.ADC/26018947).

Although the MinKNOW software conducted an initial quality filter and binning by barcode, we performed additional quality control filtering to ensure that we could screen out misbinning (either due to barcode crosstalk or the higher native error rate) and thus reduce erroneous findings. For eight barcode blanks (not included in the library preparation), an average of 970 ± 151 reads were assigned to barcodes by the software. Examination of the sequencing summary file for these reads showed most assignments were based on short barcode matches, often on one end and of relatively low identity. The secondary filtering conducted as described above (requiring a barcode match of at least 37 bases and higher identity) removed over 90% of misassigned reads to an average of 93 ± 28 reads remaining. This represents misassignment of less than 0.078% of the average number of reads per barcode used (120,036 reads/barcode).

When this same filtering process was applied to all the barcoded libraries present in the study, an average of 39.66 ± 8.29% of reads were retained.

### *Ae. aegypti* host usage in Houston, TX and Phoenix, AZ

One purpose of this study was to utilize a third-generation sequencing method for BMA, allowing us to disambiguate multiple blood meals. This would be difficult to do using current methods that rely on Sanger sequencing or methods like ELISA that require host-specific antibodies. While ELISA-based methods can detect multiple feedings (if antibodies for each specific host are used), deep sequencing based methods have shown promise to easily detect multiple blood meals as host identification is made on individual reads [20, 25]. In this study, we identified hosts from 81% (25/31) of the blood fed samples from Harris County, TX (Fig 2). Of the 25 samples with a blood meal identification, a single host was identified for 22 samples and 2 hosts were indicated for 3 samples.

**Figure 2:**
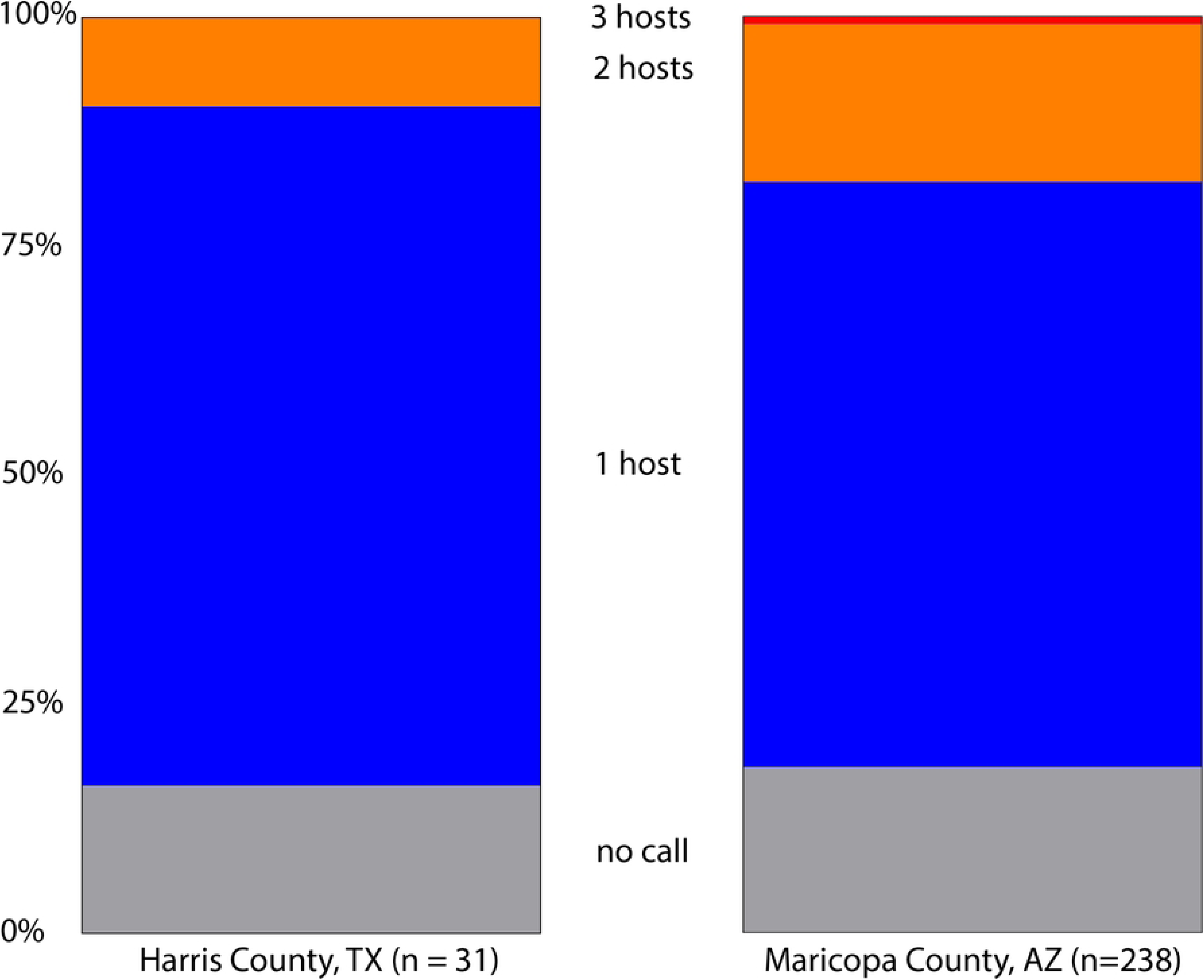
Host identification success and host multiplicity in blood fed *Aedes aegypti* collected from Harris County, Texas in 2019 and Maricopa County, Arizona in 2021 using a Nanopore based sequencing method. Minimum threshold for host identification is defined as at least two sequences with alignment length >400 bases and identity to the host *cytochrome oxidase I* gene of greater than 97.5%. Initial filtering required barcodes at each end >37 bases in length and >95% identity. Note that three hosts were identified in two samples from Maricopa County, Arizona.

Initial visual screening of Maricopa County samples identified 238 *Ae. aegypti* with evidence of a blood meal. Using this method, we determined a blood meal for 82% (195/238) of these samples (Fig 2). As with the samples from Texas, most samples indicated feeding on a single host, however 43 samples contained DNA from multiple hosts. In 2 cases, we identified the presence of DNA from 3 distinct hosts. Multiple host feeding has been observed in populations of *Ae. aegypti* from several locations [20, 24]. Statistical comparisons of Harris and Maricopa County host multiplicity were not statistically different (Fishers Exact test; *P* = 0.1791).

### Host usage in *Ae. aegypti* from Harris County, TX

BLAST results identifying *Ae. aegypti* hosts were obtained from 81% (25/31) of mosquitoes screened from Harris County. Sequence data showed that these *Ae. aegypti* were frequently feeding on felines (44%), humans (28%), and canines (20%) (Figure 3A). Three mosquito blood meals contained DNA from multiple hosts: 2 indicated feeding on human and feline, and one indicated human and canine. In these samples, reads assigned to human DNA were more abundant than the reads assigned to feline or canine. Previous BMA studies have shown a similar limited host range [12, 18]. While we only detected feeding on humans and common pets in these Harris County samples, we consider that the host range may be larger than we observed [22, 26]. While we are confident that our results are accurate, our host range findings may have been limited by the relatively small sample size.

**Figure 3:**
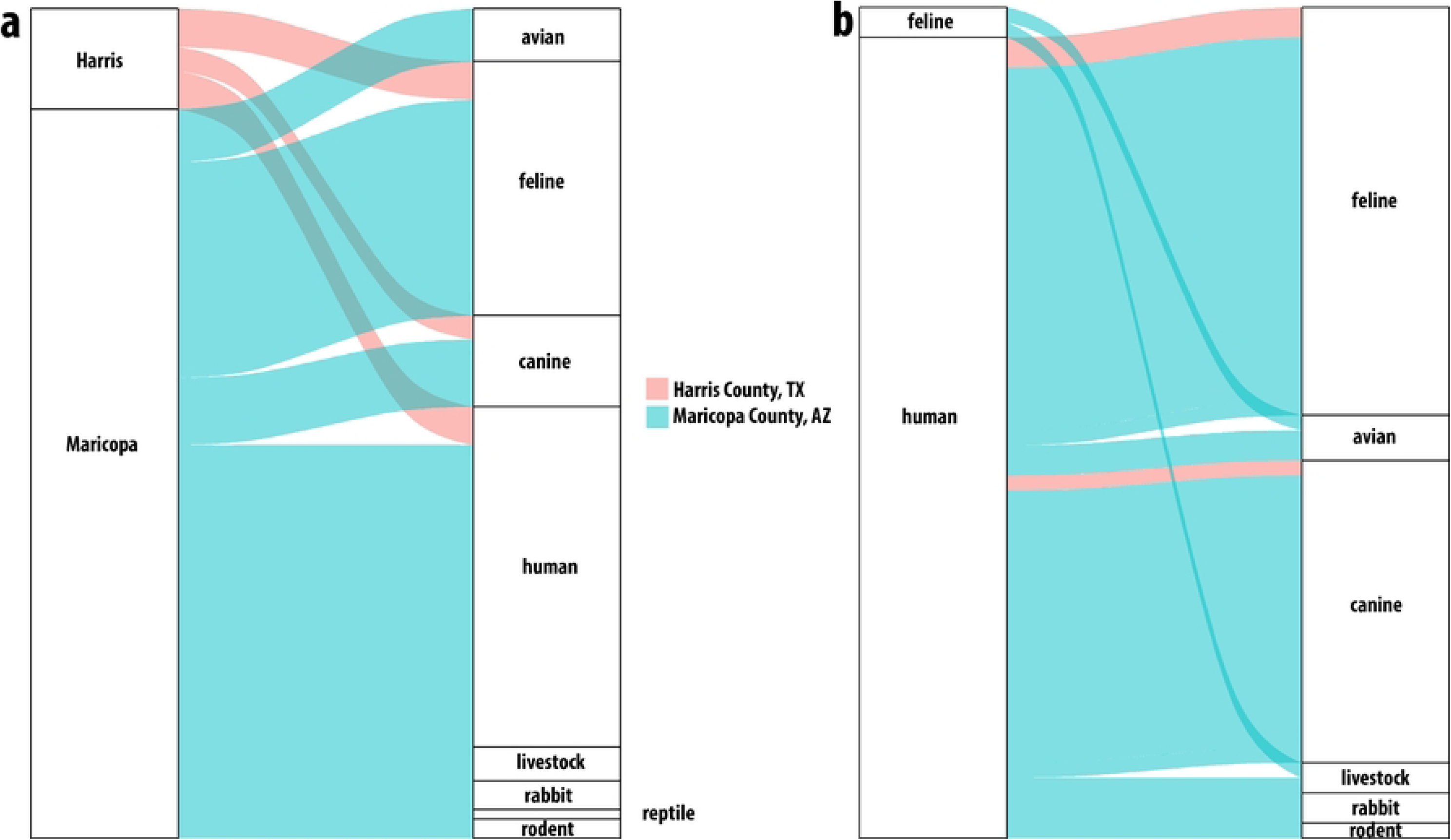
**Host usage in Harris County, Texas and Maricopa County, Arizona**. Alluvial plot showing (a) single host identifications. All samples from Harris County identified feline, human or canine hosts while Maricopa County results indicated a broader host range including birds, livestock and even reptiles. Multiple host usage (b) was observed from both locations. In Harris County, the combinations were human with feline or canine while in Maricopa County, the combinations were much broader.

### Host usage in Maricopa County *Ae. aegypti*

Detected host usage in Maricopa County was broader than that observed in Harris County (Figure 3A & 3B) and shows, as with Harris County, that local *Ae. aegypti* are not as strongly anthropophilic as might be expected based on previous studies [4, 5, 11, 12, 25, 36, 37]. While humans, felines, and canines represented more than half of the host identifications in Maricopa County, the next most frequent host group was birds, with numerous detections of feeding on chickens and several identifications of feeding on owls including the great horned owl, *Bubo virginianus*, and the long-eared owl, *Asio otus*. We also identified feeding on members of the *Zenaida* genus of doves. We identified one detection of feeding on a hawk of the *Accipiter* genus, of which *Accipiter atricapillus* is present in the Phoenix area[38].

We also detected feeding on rodents, several common livestock animals, and rabbits, which have been identified as hosts in other studies of *Ae. aegypti* feeding. We also detected feeding on ectotherms, with one identification of the desert tortoise, *Gopherus agassizii* and one identification of feeding on the ornate tree lizard, *Urosaurus ornatus*. We could only find reports of *Ae. aegypti* feeding on ectotherms from 2 areas; studies of host usage in *Ae. aegypti formosus* in Kenya and a recent study from northern Mexico, presumably from relatively similar ecological conditions as the samples tested here, that showed evidence of a tortoise blood meal[9, 20, 25].

The results from Maricopa County also indicate that taking multiple blood meals was common in these *Ae. aegypti.* Twenty-eight of the 40 multiple host samples had human and also feline or canine, likely organisms domiciled together and thus represent a convenient source of blood. Some of the other dual host results were more surprising; one sample had both feline and bird DNA present, and another had both feline and livestock DNA present. Notably, we detected two samples with evidence of three host feedings. While this was uncommon, it is not novel.

Three and even four host feeding have been previously reported in field collected *Ae. aegypti* and in laboratory studies[12, 39].

## Conclusions

Across both counties examined in the study, human DNA was found in approximately 50% of all the *Ae. aegypti* blood meals identified, followed in decreasing amounts by feline and canine. Single host blood meals for feline, canine and human did not differ between the counties (Fisher’s Exact test; *P* = 0.4216). We did identify a broader host range in the samples from Maricopa County where we detected feeding on a variety of other mammals and even reptiles.

While this broad range of hosts is interesting, we are hesitant to ascribe any real significance when we consider this in context of other BMA studies (Fig 4) in *Ae. aegypti*. Previous studies found that both host multiplicity and host range vary widely; some studies show humans are hosts for *Ae. aegypti* exclusively, while others show little to no feeding on humans (Fig 4).

**Figure 4:**
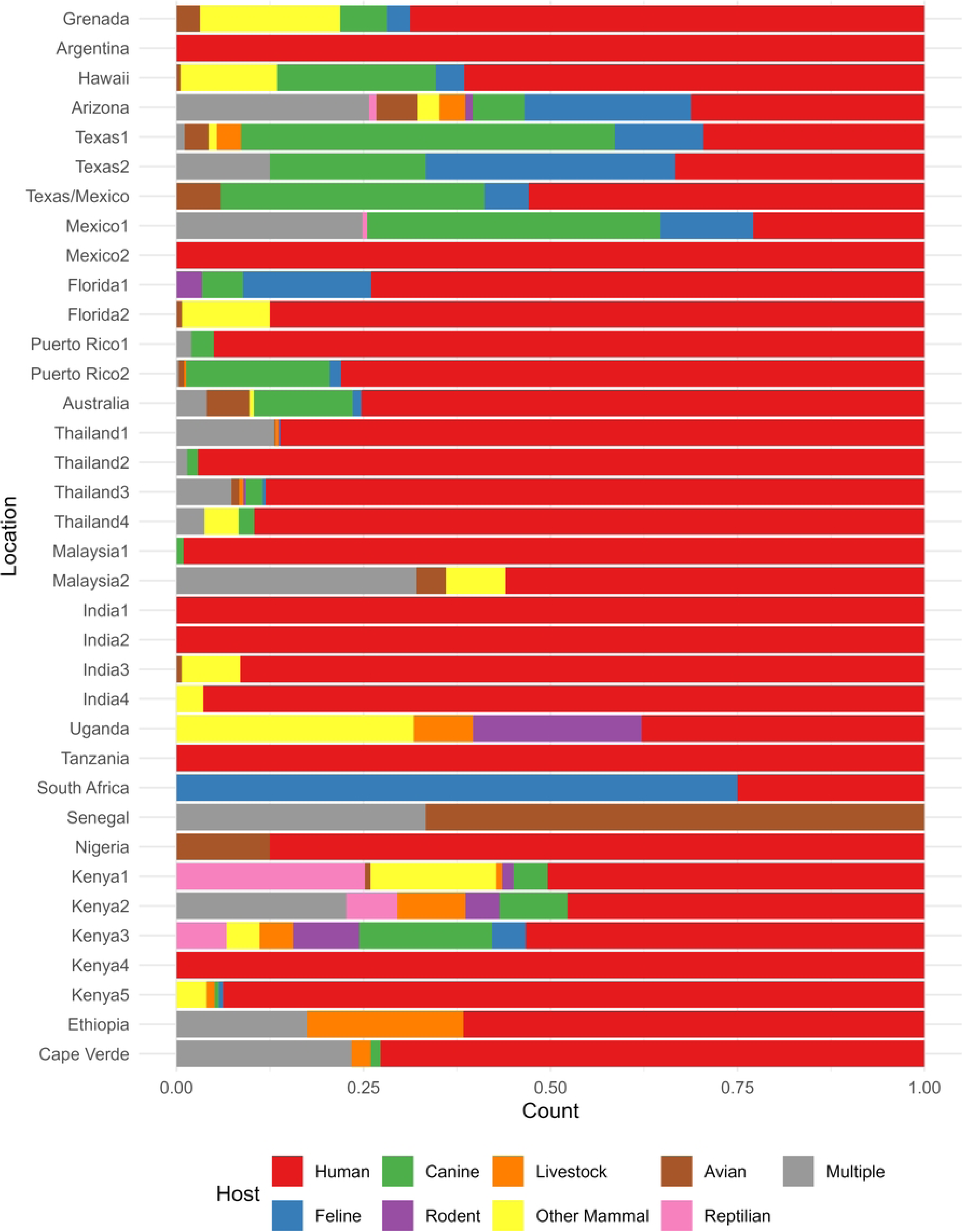
Summary of host usage studies in *Aedes aegypti*. Values are expressed as percentage and the source of the study is listed at the right. Results from this study from Harris County, TX and Maricopa County, AZ are denoted as Texas2 and Arizona respectively. Data sources are: Grenada [42], Argentina [43], Hawaii [11], Texas1 [22], Texas/Mexico [26], Mexico1 [25], Mexico2 [44], Florida1 [18], Florida2 [17], Puerto Rico1 [36], Puerto Rico2 [14], Australia [23], Thailand1 [12], Thailand2 [45], Thailand3 [5], Thailand4 [24], Malaysia1 [10], Malaysia2 [46], India1 [10], India2 [10], India3 [47], India4 [48], Uganda [9], Tanzania [49], South Africa [50], Senegal [51], Nigeria [7, 22], Kenya1 [9], Kenya2 [52], Kenya3 [52], Kenya4 [8], Kenya5 [53], Ethiopia [54], Cape Verde [55]

Similarly, some studies indicate one host, a few hosts, or a wide variety of hosts (Fig 4). These variations could be due to limitations of method (limited anti-host antibodies, inability to parse multiple overlapping Sanger electropherograms), limitations in host usage due to collection site selection (inside houses versus green space collections) or limitations in analysis (missing species-specific sequences in databases). The differences we observed may be the result of differences in host availability and convenience rather than actual differences due to preference. *Aedes aegypti* have demonstrated high adaptability by movement from a native sylvatic environment to an invasive in urban environment[40, 41]. Along with this habitat shift, it is reasonable to expect host usage would shift as host availability changes. Considering the literature, there is evidence that *Ae. aegypti* host range is determined by convenience rather than strong preference.

The detection of a diverse range of hosts in Maricopa County may also provide insight into ecological strategies implemented by *Ae. aegypti* to survive inhospitable environments.

Finding evidence of feeding on livestock, birds, rodents, and even reptiles may indicate *Ae. aegypti* are sheltering in cryptic outdoor locations like livestock barns, chicken coops, burrows, and in vegetation away from direct exposure to daytime temperatures above 40 °C and with low humidity. Subsequent collections by overnight CO_2_ baited EVS traps would indicate they may have shifted feeding activity to include periods when more suitable environmental conditions are available. Notably, six samples were positive for equine blood, and these were collected from three distinct sites. Visual examination of satellite imagery indicates evidence of horses within 150 m of the trap location for two of the three sites.

Considering all the *Ae. aegypti* host range studies (Fig 4), there is evidence that multiple host feeding is relatively common in *Ae. aegypti* and is likely underreported as both antibody-based and DNA-based BMA methods are very limited in their ability to detect multiple hosts of the same species without additional testing. Local dengue transmission in urban cycles without the involvement of zoonotic hosts and studies that indicated blood from multiple individuals within households also indicates that multiple feedings on the same species is common [39, 56]. We suggest that resampling previously tested areas with high levels of human feeding and testing these with modern deep sequencing based methods be done to determine whether *Ae. aegypti* have become regionally adapted to a restricted host range, those blood sources available to them, or if host range plasticity is widespread. This information has public health implications as a high diversity of hosts is potentially beneficial in lowering vector borne disease risk to humans.

Next generation sequencing and TGS based methods of BMA have several advantages over traditional methods and are increasingly common [19, 20, 31, 57, 58]. Like Sanger-based BMA, the ability to identify unexpected hosts improves as the number of species-specific *COI* sequences available in public databases continues to increase. In this study we used only about 500,000 well curated chordate sequences from the approximately 9 million species-specific *COI* sequences available in the latest version of the BOLD database.

Mass sequencing methods for BMA do not require specific sample types, such as excised blood-filled abdomens [16]. The methods developed for this study used DNA isolated from whole individuals, such as is commonly available to mosquito control programs. This provides more utility for samples used during routine pathogen testing and insecticide target-site resistance studies, which are commonly the primary efforts of *Ae. aegypti* surveillance. Often, the presence of a blood meal is not detected until samples have been homogenized, at which point the host and mosquito DNA have mixed and the host DNA is highly diluted, and host amplicons are rare relative to the quantity of mosquito amplicons after PCR. The TGS method described in this study is very sensitive to low host amplicon sequences among many mosquito sequences as we could detect host reads at less than 1:10,000.

The use of TGS based BMA also allows for the resolution of multiple hosts, which is not easily possible using the traditional Sanger-based method as multiple host DNA signals result in multiple ambiguous peaks in the electropherograms. Unintended amplification of mosquito *COI* by putatively vertebrate specific primers as we saw in our study (see supplemental information file 1) would result in ambiguous Sanger results but can be screened out bioinformatically since host and mosquito sequence data are easily parsed. While amplification of the mosquito DNA is detrimental for Sanger based BMA analysis, using a whole mosquito homogenate for deep sequencing is a positive as it allows for molecular identification of the mosquito species, thereby decreasing the need for morphological identification and having to discard damaged samples.

The use of mass sequencing data for BMA does require consideration of and mitigation of the limitations that are common to any database comparison method. The ability to make identifications is limited by the expansiveness of the database used for comparison. Outdated or poorly curated reference databases and preconceived assumptions of blood feeding behavior (such as that female mosquitoes only take blood meals from vertebrates) will exclude potential BLAST matches [2]. Also, careful determination of thresholds for quality filtering need to come into play to determine whether an identification goes to genus or to species. The detection of hawk as a host in Maricopa County is a good example: our database identified the reads as from *Accipiter (Astur) gundlachi*, a hawk endemic to Cuba. Closer examination of the data shows two limitations of a database comparison method. First, *Accipiter atricapillus,* which is present in Arizona, is not in the database. Second, as a matching method, the sequences for *Accipiter (Astur) gundlachi*, which were in the database, were the best match possible but at levels just above threshold. This result is not likely based the known range of *Accipiter (Astur) gundlachi*, so we must exercise discretion when determining species and genus level identification.

Data obtained for BMA using nanopore technology must be carefully screened for erroneous sequences using several layers of quality control filtering and the development of analysis pipelines that can remove misassignments. These include filtering to remove errors introduced in the process of sequencing and initial binning as well as functional filtering so that only the highest quality full amplicon sequences are used for host assignments.

Our observations from these two locations have shown that trying to generalize host usage of *Ae. aegypti* is complicated and there are many aspects that need consideration.

Researchers have historically tried to label feeding patterns of *Ae. aegypti* in hopes to improve control. Our analysis was able to highlight that in locations with environmental pressures, *Ae. aegypti* will adapt and utilize the hosts that are available in cryptic habitats and lower temperature niches to ensure species survival. Additionally, knowing what *Ae. aegypti* feed on could lead to a targeted, more informed strategy for control. For example, a blood meal analysis determining high utilization of canines by *Ae. aegypti* may indicate that treatment for mosquitoes near dog parks, kennels, etc. would be beneficial. Integrating this BMA method enables simultaneous detection of different targets of interest and will excel sapience of epidemiology of vector-borne diseases.

